# Impact of rare and common genetic variation in the Interleukin-1 pathway on human cytokine responses

**DOI:** 10.1101/2020.02.14.949602

**Authors:** Rosanne C. van Deuren, Peer Arts, Giulio Cavalli, Martin Jaeger, Marloes Steehouwer, Maartje van de Vorst, Christian Gilissen, Leo A.B. Joosten, Charles A. Dinarello, Musa M. Mhlanga, Vinod Kumar, Mihai G. Netea, Frank L. van de Veerdonk, Alexander Hoischen

## Abstract

**Background:** Interleukin(IL)-1 signaling is of major importance in human innate cytokine responses. Common variants in related genes have been linked to various inflammation-mediated diseases and stimulation-induced cytokine responses, but the role of rare variants remains to be elucidated.

**Methods:** In this study, we characterize the role of rare and common genetic variation, as identified by molecular inversion probe-based sequencing, in 48 genes related to the IL-1 pathway. Using a systems biology approach, we examined the inter-individual variability of *in vitro* stimulation-specific human cytokine responses from 463 healthy individuals of the Human Functional Genomics Project and assessed the role of rare and common genetic variants, separately and combined, by means of the Sequence Kernel Association Test.

**Results:** We identified strong associations for rare genetic variants in *NCF4* (_adj_P=7.2E^−05^) and *CASP1* (_adj_P=3.0E^−05^) with IL-6 production in response to PHA and LPS stimulation, respectively. In addition, common variants in *IL36A* and *IL38* were associated to both *C. albicans*-induced IL-1β (*IL36A* _adj_P=0.0442; *IL38* _adj_P=0.0092) and IL-6 production (*IL36A* _adj_P=0.0037; *IL38* _adj_P=0.0082), an effect that was stronger at the subpathway level both for IL-1β (_adj_P=0.0017) and IL-6 (_adj_P=1.8E^−04^). The common variant signature for the IL-1β and IL-6 response to *C. albicans* was confirmed by an association with all anti-inflammatory genes (_adj_P=1.87E^−03^ and _adj_P=5.75E^−04^), and we validated this finding for non-coding common variants. Lastly, we identified a burden of rare variants in pro-inflammatory genes and LPS-induced IL-6 production (_adj_P=2.42E^−04^), and a new role for anti-inflammatory rare variants on *S. aureus*-stimulated IL-6 production (_adj_P=6.71E^−03^).

**Conclusions:** In conclusion, we show that both common and rare genetic variation in genes of the IL-1 pathway, separately and combined, differentially influence *in vitro* cytokine responses to various stimuli in healthy individuals. This study therefore accentuates potential mechanisms that are translatable into new hypothesis-driven characterization of common and rare variant involvement in a wide variety of inflammatory and immunological mechanisms and diseases.

## BACKGROUND

The innate immune system is our first line of defense against invading pathogens such as viruses, bacteria, fungi and parasites. It consists of a variety of cell populations (*e.g*. myeloid cells, natural killer cells, innate lymphocytes) and soluble mediators (*e.g*. complement), each fulfilling a well-defined function. Collectively, these components shape innate immune responses. The induction of inflammation in response to *e.g*. infection is a naturally occurring and necessary process. Notwithstanding, both stimulatory and inhibitory mechanisms are required to induce effective elimination of pathogens, as well as to limit collateral damage to the tissues, *i.e*. prevent auto-inflammation(1).

The Interleukin-1 (IL-1) family of cytokines and receptors plays a major role in the induction and regulation of host defense and inflammation(2). The IL-1 family comprises pro-inflammatory cytokines (*e.g*. IL-1α/β, IL-36α/β/γ), anti-inflammatory cytokines (*e.g*. IL-37, IL-38), activating receptors (*e.g*. IL1-R1, IL-36R), decoy receptors (*e.g*. IL-1R2, IL-18BP), and additional regulators, kinases and phosphatases that altogether are responsible for the IL-1-mediated response(3). Next to core IL-1 family effectors, members of the inflammasome and autophagy pathway are important contributors to the regulation of IL-1-induced inflammation. For instance, activation of the inflammasome allows for cleavage and activation of caspase-1, with subsequent activation and release of pro-inflammatory cytokines IL-1β and IL-18. Conversely, autophagy is an important process involved in cell homeostasis, but is also able to directly inhibit the inflammatory response by removing inflammasome components and damaged mitochondria(4).

Defects in IL-1 pathway signaling and its specific members have been linked to various inflammation-mediated diseases(2, 5). Pro-inflammatory members of the IL-1 family, *e.g*. IL-1β and IL-18, play an important role in a variety of (auto-)inflammatory or immune diseases. For instance, a loss of balance in processing and secretion of IL-1β is an important feature of chronic inflammatory conditions like gout, systemic-onset juvenile idiopathic arthritis (sJIA), adult-onset Still’s disease (AoSD) and osteoarthritis(5). Moreover, IL-18 plasma concentrations are moderately elevated in systemic lupus erythematodes (SLE) and in rheumatoid arthritis (RA), while being severely increased in patients with macrophage activation syndrome (MAS)(5). Circulating concentrations of anti-inflammatory members of the IL-1 family are characteristically increased in numerous inflammatory conditions. Specifically, plasma concentrations of IL-1Ra, the anti-inflammatory competitor of IL-1α and IL-1β encoded by the *IL1RN* gene, are systemically increased in *i.e*. sepsis, Crohn’s disease and ulcerative colitis, and locally increased in the joints from patients with RA and osteoarthritis(6). The IL-1 type 2 receptor, a decoy receptor that binds IL-1 but does not induce an intracellular pro-inflammatory signal, has also been found increased in sepsis and has potential as a candidate biomarker in patients with acute respiratory distress syndrome(6).

For a number of (auto-)inflammatory diseases, the clinical presentation clearly points towards dysregulated activity of the IL-1 pathway. A targeted search based on clinical presentation for genetic defects in genes related to the IL-1 pathway, has therefore been proven most useful in patients with CAPS (cryopyrin associated periodic syndromes) and DIRA (deficiency of IL-1 receptor antagonist) with the identification of mutations in *NLRP3* and *IL1RN* respectively(5, 7). On the other hand, even though Adult-onset Still’s Disease (AoSD), Behcet’s and Schnitzler disease share clinical similarities with CAPS, for those no single causal genetic defect has been identified to date, despite the fact that subsets of patients have presented with mutations in related genes indicating that the likely genetic basis is close(7). In addition to rare variants, a number of Genome Wide Association Studies (GWAS) have identified common variants in the IL-1 pathway to be associated with inflammatory diseases and stimulation-induced cytokine responses(8–11). However, most studies so far have investigated rare and common variants separately. Considering increasing evidence that variability in phenotypic presentation can be caused by an interplay between variants of variable frequencies(12–14), or aggregation of genetic variants over genes underlying dysregulated biological mechanisms(15), or even over genes that are more distantly involved(16), we hypothesize that studies on the genetic basis of inflammatory diseases might also benefit from this concept.

Immune responses are highly variable between individuals, and the broad spectrum of this inter-individual variability ranges from healthy, *e.g*. protection against infection or trauma by inflammatory responses, to diseased individuals, *e.g*. sepsis or auto-inflammatory syndromes caused by excessive inflammation. For this reason, in the past few decades various studies have focused on determining the genetic variation that contributes to the inter-individual variability in immune responses(17–22). In brief, these studies assess the separate and shared contribution of host and environmental factors to an immunological response after a specific stimulus, yet all conclude that a considerable percentage of immune response variation between individuals remains unexplained. One important shortcoming is that most studies to date have focused on common genetic variants, while the impact of rare or private variants remains poorly or not at all understood. With recent advancements in sequencing technologies, the ability to study the role of rare variants has remarkably improved, and its value has been proven in several studies. For instance, increasing evidence is showing that the combined effect of common and rare variants could partially explain the missing heritability problem in complex diseases(12–14, 23, 24). The relatively small-to-moderate effects of common variants can be significantly modified by the presence or absence of (multiple) rare variants(25). Molecular Inversion Probe (MIP) based re-sequencing is a targeted sequencing technology that can identify common, low-frequency, rare and even private variants within a region of interest, and as such allows to cost-effectively study intermediate sized gene panels (*e.g*. 50 genes) in relatively large cohorts(26–29).

Considering the importance of the IL-1 pathway for inflammation and innate immune responses, we aimed to identify and characterize rare and common genetic variants in 48 genes related to the IL-1 pathway mediated immune response, and determine their impact on the inter-individual variability of cytokine responses in healthy individuals. A complete overview of the study workflow can be found in **Figure 1**.

**Figure 1.**
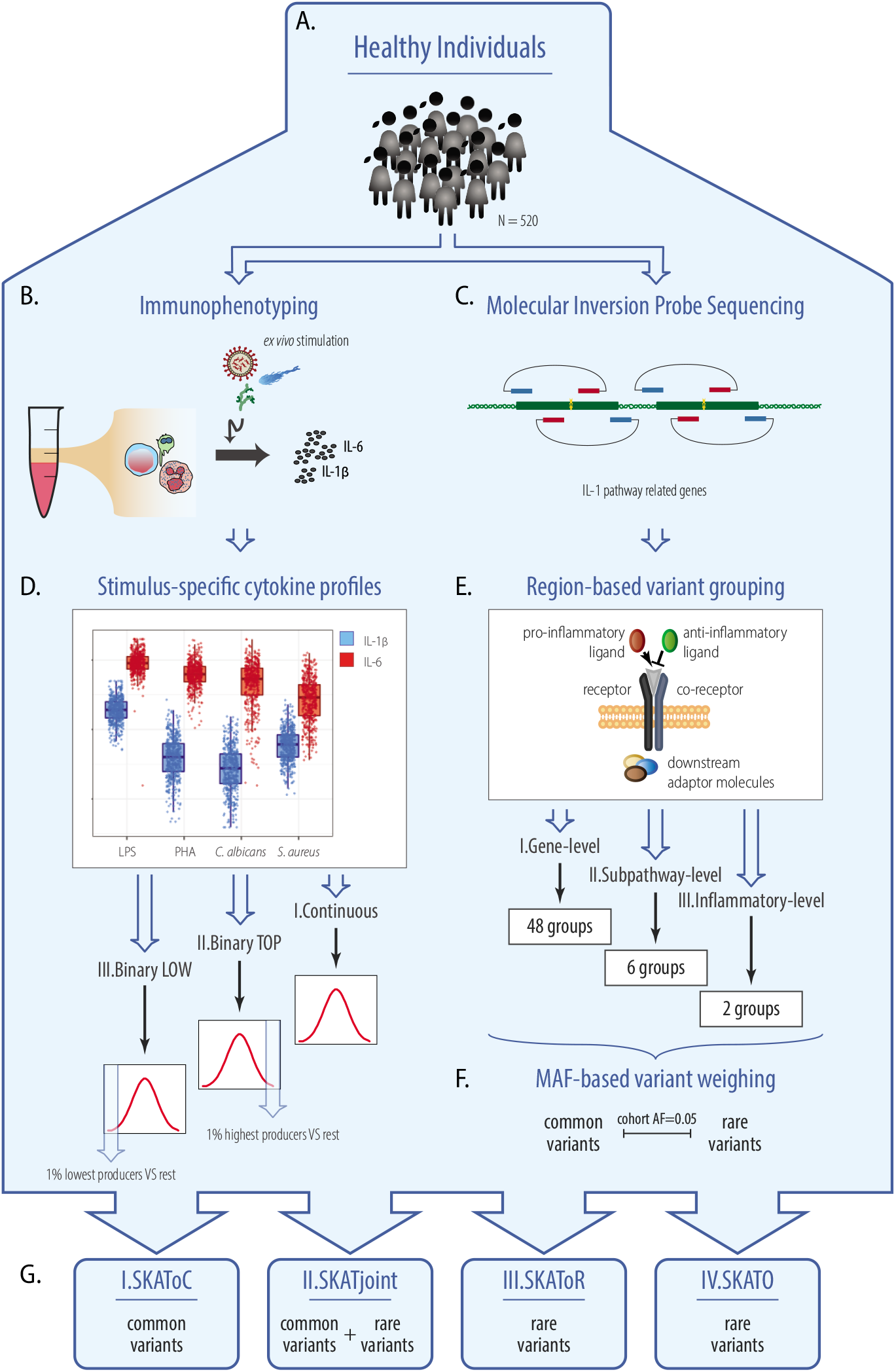
Flowchart of the study workflow method. Figure orientation from top to bottom. Blood was extracted from 520 healthy individuals **(A)** on which extensive immunophenotyping was performed **(B)** and simultaneously Moleclar Inversion Probe sequencing data was produced from coding regions of 48 Interleukin-1 pathway genes **(C)**. The resulting cytokine production after stimulation was measured and used for analysis **(D)**: I.Continuous, using log-transformed IL-1β and IL-6 cytokine production upon stimulation; II. Binary TOP, testing the 1% highest IL-1β and IL-6 cytokine producers versus the rest; III. Binary LOW, testing the 1% lowest IL-1β and IL-6 cytokine producers versus the rest. Similarly, the identified variants were grouped over three different regions into sets based on gene-encoded protein function **(E)**: I.Gene-level, with 48 gene-groups; II.Subpathway-level, with 6 subpathway-groups; and III.Inflammatory-level, with 2 inflammatory-groups. Variants within each set were appropriately weighed based on Minor Allele Frequency (MAF), and common- and rare variants were classified based on cohort Allele Frequency (AF) threshold of 0.05 **(F)**. Finally, variant analysis was performed by the Sequence Kernel Association Test (SKAT): I.SKAToC, SKAT with only common variants; II.SKATjoint, SKAT with common and rare variants; III.SKAToR, SKAT with only rare variants, and IV.SKATO; best combination of the SKAT and Burden Test with only rare variants **(G)**.

## MATERIAL AND METHODS

### Study Cohort

#### Cohort characteristics

The study was conducted using healthy individuals from the Human Functional Genomics Project (HFGP; 500FG-cohort)(30). The entire 500FG-cohort consists of 534 healthy individuals from the Netherlands (296 females and 237 males) with an age range 18-75, from which we were able to obtain DNA from 520 individuals for sequencing. For more details on cohort characteristics see previous publications on the 500FG-cohort(17, 18, 20).

#### Immunophenotyping

In this study we made use of the publicly available extensive immunophenotyping data that was generated as part of the Human Functional Genomics Project(31). Specifically, Interleukin-1ß (IL-1ß) and Interleukin-6 (IL-6) production by whole blood (consisting mainly of polymorphonuclear cells (PMNs)) from 471 individuals, stimulated with either lipopolysaccharide (LPS, 100ng/mL), phytohaemagglutinin (PHA, 10μg/mL), heat-killed *Candida albicans* (*C. albicans* 10^6^ CFU/mL) or *Staphylococcus aureus* (*S. aureus* 1 x 10^6^/mL). A detailed description of these experiments can be found elsewhere(18). In short, blood was drawn from participants and 100 μL of heparin blood was stimulated with 400 μL of stimulus, subsequently incubated for 48 hours at 37°C and 5% CO_2_ and supernatants were collected and stored in −20°C until used for ELISA.

### Sequencing

#### MIP-panel design

We sequenced all coding exons of 48 genes of the IL-1 pathway in 520 healthy individuals by Molecular Inversion Probe (MIP)-based re-sequencing. MIPs are a targeted re-sequencing technology, that allows for the identification of both common- and rare genetic variation in regions of interest. A detailed description of MIP-probe design and sequencing methods can be found elsewhere(26, 29, 32). In short, 1285 MIP-probes were designed to cover all coding exons of 48 genes related to the IL-1 pathway and sequencing was performed using the Illumina NextSeq500 system. These 48 IL-1 pathway related genes can be further functionally subclassified into six subpathways that represent a sub-mechanism or immunological cascade in the IL-1-mediated inflammatory response: IL-1 subpathway, IL-18 subpathway, IL-30s subpathway, Inflammasome, (reactive oxygen species) ROS-production, and Autophagy. An additional functional sub classification into two inflammation phenotype groups, was based on the gene-encoded protein function and its pro- or anti-inflammatory effect. A full explanation on the sub classifications can be found in **Additional File 1**.

#### Data processing

A carefully developed filtering pipeline, based on Sanger Sequencing validations, was applied to ensure high sensitivity and specificity in our final variant set. First, the reads were aligned using BWA-MEM(33) and subsequently filtered on Mapping Quality ≥ 60, no soft-clipping, properly paired and not more than five variations from the reference per read, with the exception of multi-basepair insertions and deletions. Variants were then called using the GATK unified genotyper(34), which uses a Bayesian genotype likelihood model to estimate the most likely genotypes. Rare variants (here defined as absent in dbSnp build 150 common(35), or defines as rare by our custom annotator as explained below), were further filtered on the QUAL parameter ≥ 1000 in the vcf. Additionally, the percentage of alternative alleles for each variant position using samtools mpileup(36), with maximum read depth 10000, no BAQ and a minimal base quality of 30. Homozygous rare variants required an alternative allele percentage of ≥ 90%, heterozygous rare variants required an alternative allele percentage of 25% and < 90%, and rare variants with an alternative allele percentage of < 25% were considered false positive. Samples with an average coverage depth of all MIPs ≥ 100x were included for analysis. The final variant set was annotated using our custom annotator, which makes use of several annotation sources, among others the Variant Effect Predictor (VEP) from Ensembl(37), Combined Annotation Dependent Depletion (CADD) score(38), and several population based variant databases (*e.g*. dbSnp, ExAc and gnomAD(39)) and an “inHouse” database consisting of > 25,000 clinical exomes run at the diagnostic division of the Department of Human Genetics of the Radboud University Medical Center (Radboudumc). We used within cohort allele frequencies (AFs) to separate rare and common variants, based on a common variant cut-off of ≥ 5%.

### Variant Analysis

#### Continuous Trait

A Rare Variant Burden Analysis (RVBA) was performed on the log-transformed cytokine levels by using the Sequence Kernel Association Test (SKAT)(14, 40) in R version 3.5.2. The SKAT is a kernel-based test method, that aggregates weighted individual variant-score test statistics whilst allowing variant-variant interactions and is extremely powerful when a genetic region has both protective and deleterious variants or many non-causal variants(14, 23, 24, 40, 41). The SKAT was performed over three levels of grouping: I) **gene-level**; where all variants within coding exons of a gene region were considered (**Figure 1E.I**), II) **subpathway-level**; where all variants within genes that belong to the corresponding subpathway were considered (**Figure 1E.II**), and III) **inflammatory-phenotype level**; where based on encoded protein function genes were classified as either pro-inflammatory or anti-inflammatory and all variants from genes in either groups were considered (**Figure 1E.III**). Furthermore, for each region the ‘normal’ SKAT was run to determine the effect of only common (I.SKAToC), only rare (III.SKAToR) and common and rare variants combined (II.SKATjoint), based on a cohort MAF of 5% as cut-off, using the SKAT_CommonRare function with default weights. However, since the SKAT can be less powerful than burden tests when rare variants in a set are truly causal or influencing the phenotype in the same direction(41), we additionally applied the SKAT-O algorithm to the rare variants (SKATO), appropriately weighing the variants with the same weights as described above, and extracted accompanying rho-values to assess the contribution of SKAT versus Burden Test for significant sets reflecting the proportion of bi- and unidirectionality of an association. In the case of rare- and joint tests only output based on > 1 variant was considered, and in the case of joint tests the presence of both rare and common variants in the set was an additional requirement. P-values were Bonferroni-adjusted for each previously defined test separately, based on the number of groups tested. We did not apply additional corrections for the different grouping-levels (*i.e*. gene-level, subpathway-level, inflammatory-phenotype level), nor for the different variant frequency tests (*i.e*. SKAToC, SKATjoint, SKAToR, SKATO).

#### Binary Trait

In addition to the associations of continuous cytokine levels as described above (**Figure 1D.I**), we performed a binary association analysis on outlier individuals, here defined as extreme cytokine producers. As research has shown that individuals with outlier expression patterns are likely to be enriched in nearby rare variants(42–45), we hypothesized that outlier individuals in terms of cytokine production could similarly be enriched in rare variants in specific genes, thereby favoring the identification of stimulus-specific mechanisms. For this purpose, we defined for each cytokine-stimulus the 1% extreme cytokine producers (rounded up, so generally ±5 individuals), which categorized the individuals in two groups that were subjected to binary trait association. Specifically, for each cytokine-stimulus combination the SKAToR was applied twice: 1) 1% highest cytokine producers versus all other individuals (_TOP_BT, **Figure 1D.II**), and 2) 1% lowest cytokine producers versus all other individuals (_LOW_BT; **Figure 1D.III**). In two cases, *C. albicans-* induced IL-1ß production low-producers and LPS-induced IL-6 production high-producers, no distinctive categories could be created due to equal cytokine measurements at the 1% cut-off, and as such the groups were extended to 7 and 9 respectively. We followed up on associations based on >1 variant, that were nominally significant (unadjusted P-values) in the continuous associations and recurrent in either _TOP_BT or _LOW_BT.

#### Follow Up

In order to give meaning to our detected associations, we extracted the residual (corrected for covariates age and gender) cytokine production from the SKAT null-model and correlated those to the genotype categories, where applicable. For all plots, correlations and cytokine levels mentioned from hereon, the cytokine production therefore has been corrected for age and gender. In the case of set-based common variant associations we correlated cytokine production to the three separate genotype categories; homozygous reference, heterozygous, homozygous alternative, whereas in the case of set-based rare variant associations we correlated cytokine production to only two categories; absence or presence of one of the rare variants in the set.

Additionally, considering accumulating evidence for a role of non-coding genetic variation in health and disease(46, 47), we followed-up on common variant associations by using the publicly available genotype data from the FG500-cohort, generated with the commercially available SNP chip Illumina HumanOmniExpressExome-8v1.0 (for further details we refer to previously published work(18, 48)). We extracted all common variants (based on cohort AF ≥ 5%) within NCBI RefSeq ‘Whole Gene’ gene regions and extended the start position by 50kB(49) for the following sets: *IL36A, IL38*, IL-30s subpathway, pro-inflammatory phenotype and anti-inflammatory phenotype. Variant sets were pruned for Linkage Disequilibrium (LD) based within cohort metrics and the commonly used R^2^ cut-off of >0.8, by means of the snpStats package in R. The final pruned set of variants, termed tagSnps, were subjected to the same SKAT with default weights, to test for association with continuous IL-1ß (N=428) and IL-6 (N=425) cytokine production. Finally, for the purpose of correlating significant non-coding common variant sets to cytokine levels, we calculated an allelic score for all variants in the set. An allelic score is a way to collapse multidimensional genetic data associated with a risk factor into a single variable(50). We slightly adapted the allelic score calculation to our SKAT-based test results, into a weighted (based on AF-based Beta.Weights function from SKAT package), directional (based on increasing or decreasing cytokine production over the genotype categories) allelic score. Specifically, we inferred the direction of each variant in a set, and combined this with the manually computed variant weight, by negating the weight only for variants with decreasing cytokine production over the genotype categories. Common variant genotypes were converted to dosages and multiplied by their directional weight, which was summed up to an allelic score per set of variants. The weighted, directional allelic score was plotted versus the cytokine production, and a linear model equation and R^2^ metrics were extracted. In addition, to evaluate the individual contribution of non-coding common variants in a set, we computed per SNP linear models using *C. albicans-induced* residual cytokine production as the criterion variable and the SNP in question as predictor variable. The individual SNP effect estimates (or Beta-estimates) were organized by direction and annotated based on their significance. For the purpose of prioritizing one SNP for investigating epigenetic effects we computed the same linear models now using the log-transformed cytokine production as criterion variable and next to the SNP also age and gender as covariates. The predictive capacity of the linear models, as reflected by the model p-value, was used as a measure for impact of a specific SNP on cytokine production and as such prioritized rs1562305 for more in-depth follow-up. Epigenetic effects of rs1562305 were studied using a Hi-C dataset(51) and Pol II ChiA-PET libraries for K562 (GSM970213). Transcription Factor ChiP-seq Clusters (338 factors, 130 cell types) were obtained from ENCODE 3 Data version: Encode 3 Nov 2018.

## RESULTS

### Study Cohort

Here, we studied healthy individuals from the Human Functional Genomics Project (HFGP; FG500-cohort)(30), by making use of the publicly available demographic data and stimuli-specific *in vitro* cytokine measurements(31). The gender distribution over 463 included individuals for analysis shows a mild overrepresentation of females as compared to males (Male n=201, Female n=262), whereas the mean- and median age distribution for these groups separately is comparable (**Additional File 2A**).

*In vitro* IL-1β and IL-6 cytokine production in whole blood in response to stimulation with either 100ng/mL Lipopolysaccharide (LPS), 10μg/mL Phytohemagglutinin (PHA), heat-killed *Candida albicans* 10^6^CFU/mL (*C. albicans*) and 1×10^6^/mL *Staphylococcus aureus (S. aureus*), were likewise evenly distributed between females and males (**Additional File 2B**), and were log-transformed prior to analysis. Based on the above-mentioned distributions in combination with the fact that previous research has shown that age and gender can influence cytokine responses(17–20), both variables were included as covariates in our analyses.

### Sequencing

Molecular Inversion Probe (MIP)-based re-sequencing of all coding exons of the 48 genes in our IL-1 pathway MIP-panel generated sequencing data from 520 healthy individuals (for all MIPs see **Additional File 3**). Overlapping the sequencing data with the available immunophenotyping data we managed to obtain complete datasets from 463 individuals for analysis. The average coverage depth for these 463 individuals over all MIPs was 830x (**Additional File 4**). Five genes in our panel (*SIGIRR, PYCARD, CYBA, RAC2* and *MAP1LC3A*) were unfavorably covered (<100x average coverage of the entire coding part of the gene) for more than half of the samples, and one gene (*NCF1*) lost all coverage in our extensive quality filtering due to homology regions, and was therefore excluded from subsequent analysis (**Additional File 4**). Based on gene-encoding protein function and the immunological cascade in which they are activated, these 48 genes were classified prior to the analysis into 1) six subpathway groups: IL-1 subpathway, IL-18 subpathway, IL-30s subpathway, Inflammasome, ROS-production and Autophagy; and 2) two inflammatory-phenotype groups: pro-inflammatory, and anti-inflammatory (**Additional File 1**).

Overall, we identified 201 non-synonymous variants in the coding exon regions, out of which 35 were common and 166 were rare (based on cohort allele frequencies (AFs) using a threshold of ≥ 5% for common variants). Furthermore, out of 166 rare variants we identified 18 variants to be novel, *i.e*. never observed before in public databases. For a complete variant list see **Additional File 5**.

### Variant Analysis

The role of rare and common variants on stimuli-specific cytokine responses was assessed by a Rare Variant Burden Analysis (RVBA) using the Sequence Kernel Association Test (SKAT)(14, 40, 41). The main issue with rare variant association testing lies in the low-frequency of rare variants, and thus appropriate grouping is warranted(23, 24). We performed the SKAT using three different grouping strategies (**Figure 1E** and **Additional File 1**): I) **gene-level**, where all variants within the coding region of each gene were considered; II) **subpathway-level**, where all coding variants within genes that belong to the corresponding subpathway were considered; and III) **inflammatory-phenotype level**, where genes were classified as either pro-inflammatory or anti-inflammatory based on encoded protein function, and all variants from genes in either group were considered. Each level was assessed for the role of genetic variants on stimuli-specific IL-1β and IL-6 cytokine production, through the full spectrum of AFs (from rare- to common (AF ≥ 5%) variants separately and combined) by performing four different tests (**Figure 1G**): I.SKAToC; testing only common variants, II.SKAToR; testing only rare variants, III.SKATjoint; testing common and rare variants jointly by using a method that considers common variant effects as covariates in the rare variant test. To verify potentially missed strong unidirectional signals, the rare variants were additionally subjected to an adapted algorithm that uses a linear combination of the SKAT and Burden Test, the SKAT-O (IV.SKATO). Output from all SKATs performed in this study can be found in **Additional File 6**.

#### Association landscapes show similarities and differences in IL-1ß and IL-6 response

We created holistic heatmap overviews termed ‘association landscapes’, for the purpose of summarizing rare- and common variant associations, both on the gene and subpathway level, in an organized fashion. **Figure 2** shows these landscapes of gene- and subpathway based associations for IL-1ß (**2A**) and IL-6 (**2B**) production by whole blood. **Figure 3** shows the inflammatory-phenotype based associations for IL-1ß (**3A**) and IL-6 (**3B**) production by whole blood in classic rectangular heatmaps.

**Figure 2.**
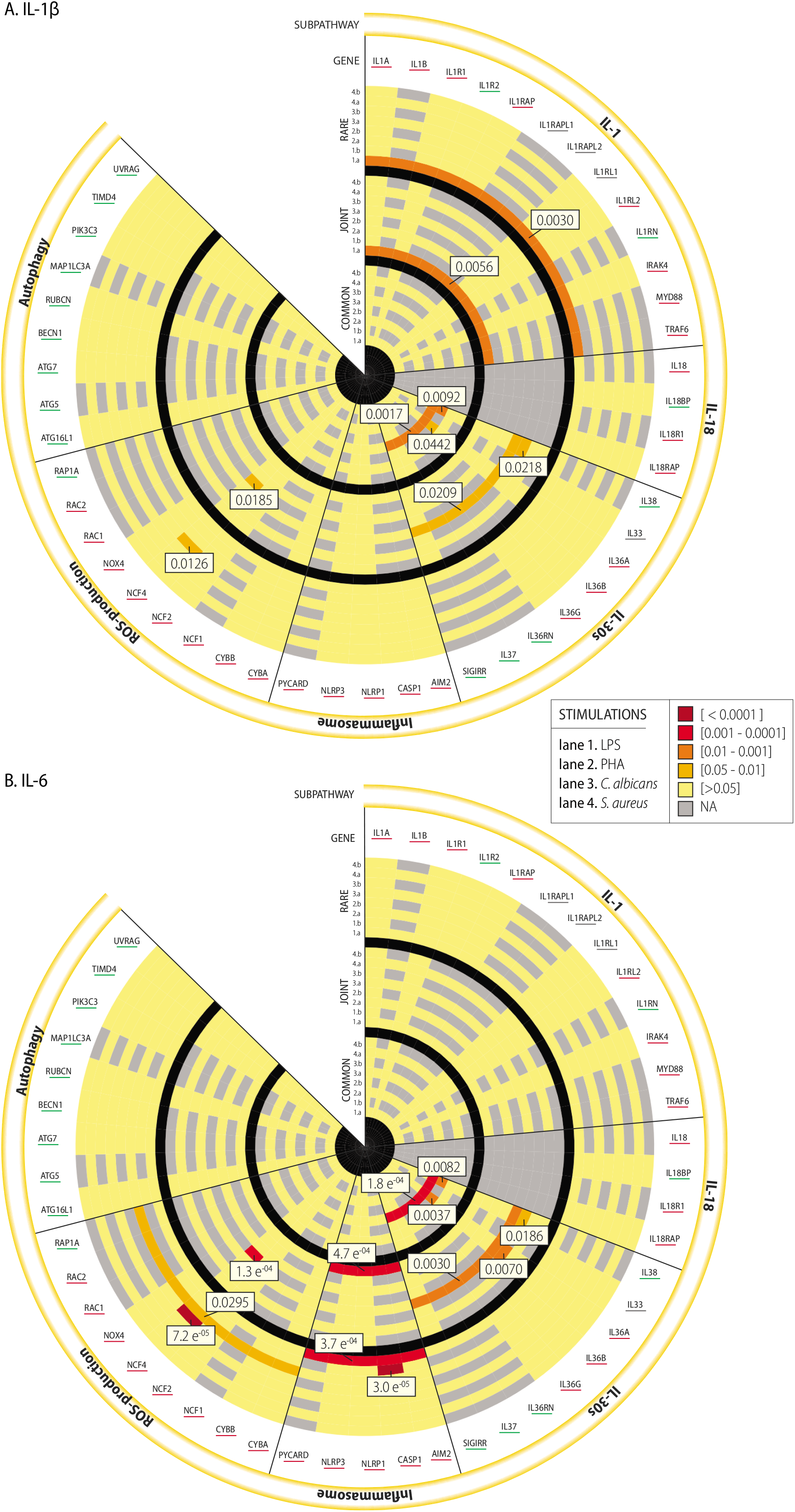
Circular heatmaps of SKAT _adj_P-values. The circular heatmaps consist of three rings separated by black lanes, representing from inner to outer the SKAToC (SKAT with only common (AF ≥ 5%) variants), SKATjoint (SKAT with common and rare variants), and SKAToR (SKAT with only rare variants) _adj_P-values with log-transformed IL-1β **(A)** and IL-6 **(B)** cytokine production respectively. Each ring consists of 8 lanes, that represent different stimuli; 1) LPS 100ng/mL, 2) PHA 10μg/mL, 3) *C. albicans* 10^6^CFU/mL, 4) *S. aureus* 1×10^6^/mL, with sub lane a) showing the subpathway-level result and sub lane b) showing the gene-level result. The genes annotated at the surface of the heatmap are underlined with a red, green or grey color, representing their classification in the inflammatory-phenotype level groups (**Figure 3**).

**Figure 3.**
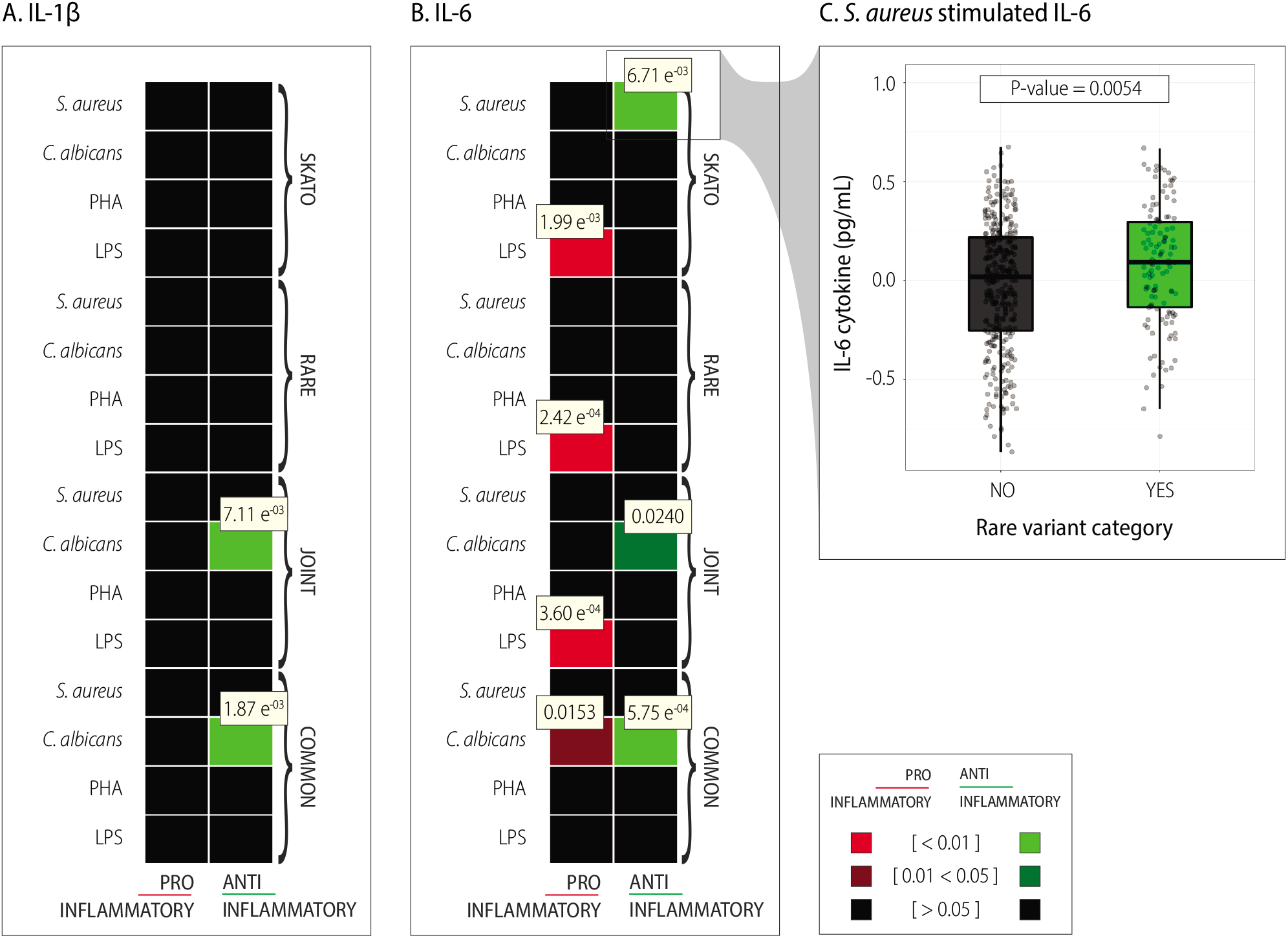
Inflammatory-phenotype level SKAT _adj_P-values. A heatmap representation of association between inflammatory-phenotype groups and IL-1β **(A)** and IL-6 **(B)** cytokine production in response to four different stimuli; LPS 100ng/mL, PHA 10μg/mL, heat-killed *C. albicans* 10^6^CFU/mL, and *S. aureus* 1×10^6^/mL. Variants in genes categorized as pro- or anti-inflammatory were subjected to four different association tests; SKAToC (common (AF ≥ 5%) variants only), SKATjoint (common and rare variants), SKAToR (rare variants only), and SKATO (combination of SKAT and Burden Test with rare variants only). **(C)** shows a boxplot of *S. aureus* stimulated residual IL-6 cytokine production (corrected for age and gender) over rare variant categories (NO=individuals without rare variant; YES=individuals carrying a rare variant) within genes of the anti-inflammatory phenotype group. Wilcoxon rank-sum test P-value reveals a significant difference between the two categories.

The **gene-level** analysis revealed significant associations between rare genetic variants in *NCF4* and both IL-1ß and IL-6 cytokine production in response to PHA stimulation (SKAToR _adj_P-value=0.0126 and 7.2E^−05^ respectively), that sustained when common variants were considered as covariates (SKATjoint _adj_P-value=0.0185 and 1.3E^−04^ for IL-1ß and IL-6 respectively). Another strong rare variant signal was observed between *CASP1* rare variants and LPS-induced IL-6 cytokine production (SKAToR _adj_P-value=3.0E^−05^). Next to this, common variants in *IL36A* and *IL38* were significantly associated to both IL-1ß and IL-6 cytokine production after *C. albicans* stimulation (*IL36A* SKAToC _adj_P-value=0.0442 and 0.0037; *IL38* SKAToC _adj_P-value=0.0092 and 0.0082). These associations substantially weakened in the joint analysis of common and rare variants, with the exception of the *IL36A* association with IL-6 that remained comparable to the rare variant result (SKATjoint _adj_P-value=0.0070).

The **subpathway-level** analysis identified a significant association between rare genetic variants of IL-1 subpathway genes combined and LPS-induced IL-1ß cytokine production (SKAToR _adj_P-value=0.0030), that maintained in the joint analysis with only a minor decrease in significance (SKATjoint _adj_P-value=0.0056). Specific to IL-6, we identified associations between rare variants in ROS-production genes after PHA stimulation (SKAToR _adj_P-value=0.0295), and rare variants in Inflammasome genes combined after LPS stimulation (SKAToR _adj_P-value=3.7E^−04^), while only the latter remained in the joint analysis (SKATjoint _adj_P-value=4.7E^−04^). Finally, common variants in IL-30s subpathway genes were significantly associated with both IL-1ß and IL-6 cytokine production in response to *C. albicans* stimulation (SKAToC _adj_P-value=0.0017 and 1.8E^−04^ respectively), both slightly weaker in the joint analysis.

The **inflammatory-phenotype level** analysis (**Figure 3**), revealed a strong association between rare variants in genes with pro-inflammatory effects and LPS-induced IL-6 cytokine production (SKAToR a_dj_P-value=2.42E^−04^ and SKATO a_dj_P-value=1.99E^−03^), that was recurrent in the joint analysis (SKATjoint _adj_P-value=3.60E^−04^). On the other hand, rare variants in anti-inflammatory genes were significantly associated with IL-6 cytokine production in response to *S. aureus* stimulation (SKATO _adj_P-value=6.71E^−03^). Moreover, we identified a role for common variants in anti-inflammatory genes in both IL-1ß and IL-6 cytokine production in response to *C. albicans* stimulation (SKAToC _adj_P-value=1.87E^−03^ and 5.75E-^04^ respectively), that in the joint analysis persisted for IL-1ß (SKATjoint _adj_P-value=7.11E^−03^), whereas in for IL-6 most of the signal was lost (SKATjoint _adj_P-value=0.0240). Finally, unique to *C. albicans-induced* IL-6 production, was an association with common variants in pro-inflammatory genes (SKAToC _adj_P-value=0.0153).

#### Gene-level rare variant associations of cytokine outliers imply cytokine-stimulus specific features

The associations presented thus far were produced by using stimulus-specific cytokine profiles of all individuals as continuous measurements. In order to uncover rare variant gene associations that may reflect strong stimulus-specific mechanisms, we supplemented these continuous trait associations (CT, **Figure 1D.I**), with extreme binary trait associations (BT). The BT associations were conducted by categorizing individuals into two groups, based on 1% highest and lowest cytokine production respectively, in response to a certain *in vitro* stimulus, from here on referred to as top-producers (_TOP_BT, **Figure 1D.II**) and low-producers (_LOW_BT, **Figure 1D.III**). Identifying recurrent rare variant associations (that is, nominally significant in CT and either _TOP_BT or _LOW_BT), allowed us to give meaning in terms of direction of the effect of our continuous association results. **Figure 4** shows recurrent gene-level rare variant association (SKAToR) P-values, separately for IL-1ß (**A**) and IL-6 (**B**).

**Figure 4.**
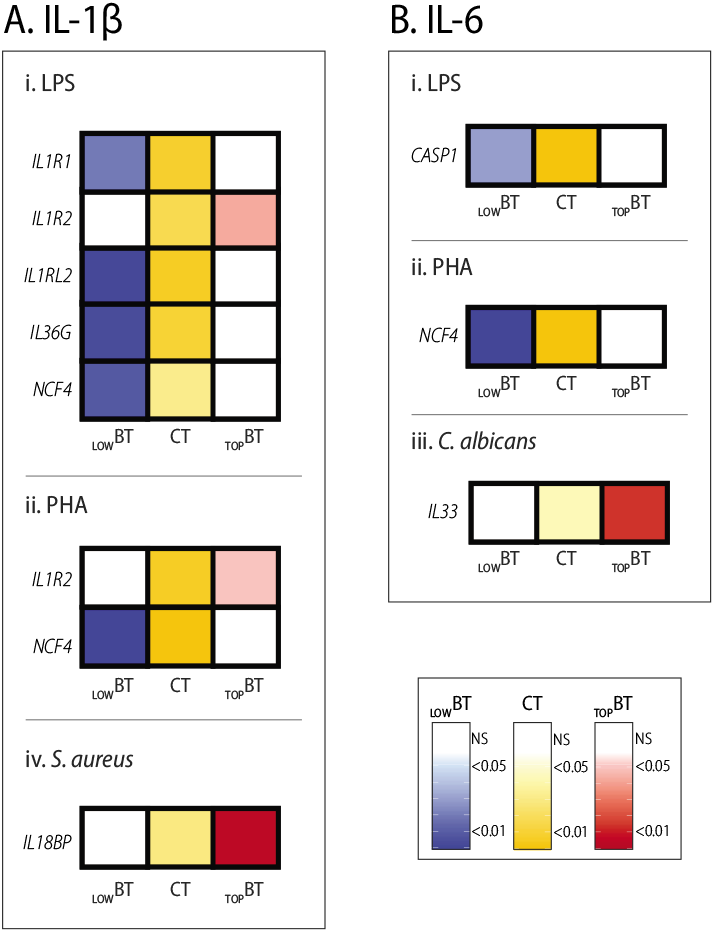
Extreme cytokine producers provide direction and identify distinct cytokine-stimulus specific features. A heatmap representation of all recurrent gene-level rare variant SKAT P-values, where recurrence is defined per cytokine-stimulus combination as a nominal significant P-value in either top- or low-producers in addition to a nominal significant P-value in the corresponding continuous association. Association between rare variants and IL-1β **(A)** and IL-6 **(B)** cytokine production in response to LPS 100ng/mL **(i.)**, PHA 10μg/mL **(ii.)**, *C. albicans* 10^6^CFU/mL, and *S. aureus* 1×10^6^/mL stimulation **(iv.)**. Annotation: _LOW_BT = SKATBinary with lowest 1% producers; CT = SKAT with log-transformed continuous cytokine producers; _TOP_BT = SKATBinary with highest 1% cytokine producers.

The analysis of extreme cytokine producers was consistent with our identified continuous association result between rare variants in *NCF4* and PHA-induced IL-1ß and IL-6 cytokine production, by the identification of a recurrent signal in the low-producers (IL-1ß _LOW_BT P-value=2.18E^−04^; IL-6 _LOW_BT P-value=5.94E^−05^), along with the association between *CASP1* and IL-6 low-producers after LPS stimulation (_LOW_BT P-value=0.0299). In addition to that, we identified three recurrent associations in IL-1ß top-producers: rare variants in *IL1R2* after LPS (P-value=0.0429) and PHA stimulation (P-value=0.0493); *IL18BP* rare variants after *S. aureus* stimulation (P-value=0.0038); and four recurrent associations in LPS-induced IL-1ß low-producers - rare variants in *NCF4* (P-value=8.27E^−03^), *IL1R1* (P-value=0.0194), *IL1RL2* (P-value=8.54E^−04^) and *IL36G* (P-value=3.03E^−03^) (**Figure 4A**). In the case of IL-6 (**Figure 4B**) we detected one additional recurrent signal, that is *IL33* rare variants and top-producers after *C. albicans* stimulation (P-value=0.0143).

#### Immunological response to C. albicans reflects a common variant signature

Our results of the SKAToC show strong common coding variant set signals over various levels of magnitude both on IL-1ß and IL-6 cytokine production after *C. albicans* stimulation, reflecting a common variant signature in this immunological response. For the separate variants in *IL36A* (rs895497) and *IL38* (rs6761276 and rs6743376), we observed that the alternative allele presented with 1) a higher frequency as compared to the ancestral allele, and 2) a higher cytokine production (residual, after correcting for co-variates age and gender). **Figure 5A** shows that for each of these variants the IL-1ß and IL-6 *in vitro* cytokine production in response to *C. albicans* stimulation decreases over the genotype categories. Complementary Wilcoxon-rank-sum tests confirmed that for all three variants homozygous reference versus homozygous alternative and heterozygous versus homozygous alternative, was significantly associated with higher IL-1ß and IL-6 cytokine production, and homozygous reference versus heterozygous only for *IL38* rs6743376 (**Figure 5A**).

**Figure 5.**
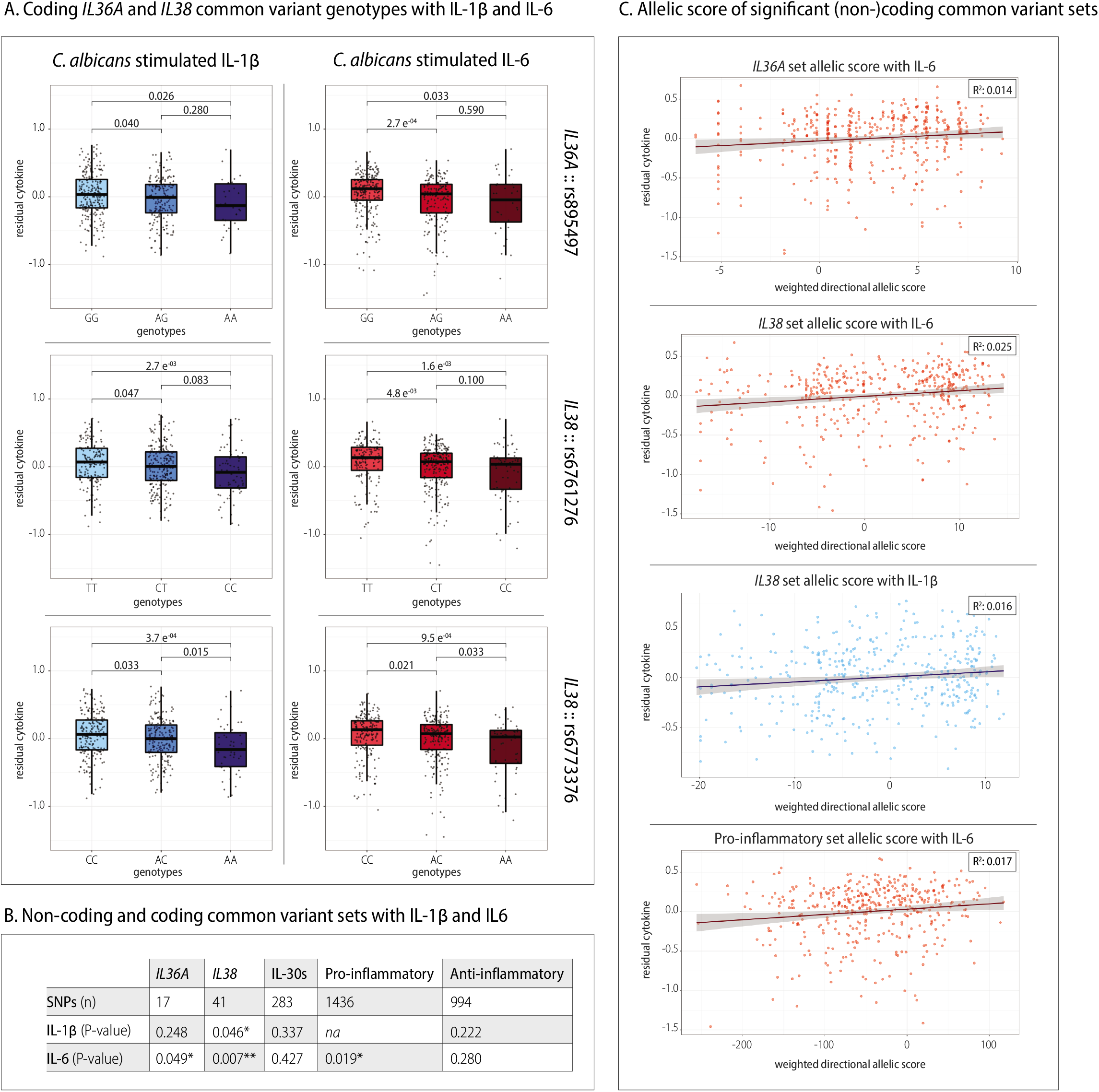
Coding and non-coding common variant set associations with *C. albicans* induced cytokine production. **(A)** shows that the residual IL-1β (left-panel in blue) and IL-6 (right-panel in red) cytokine production for coding SNPs in *IL36A* and *IL38* decreases over the genotype categories. For all plots the ancestral allele is the minor allele and thus the genotype categories are ordered from left to right: homozygous alternative (IL-1β in light-blue and IL-6 in light-red), heterozygous (IL-1β in mid-blue and IL-6 in mid-red), homozygous ancestral (IL-1β in light-blue and IL-6 in light-red). **(B)** confirms that next to coding common variants, also non-coding common variants in *IL36A, IL38* and Pro-inflammatory phenotype sets are associated with *C. albicans* induced residual IL-6 cytokine production, and *IL38* set with *C. albicans* induced residual IL-1β cytokine production. **(C)** shows the Beta-weighted, directional, allelic score in correlation with *C. albicans* induced residual IL-1β or IL-6 cytokine production. The straight line represents the linear model equation using method ‘lm’ with standard error of 0.95, and the R^2^ (measure of correlation) is displayed in the plot. Annotation: * = P-value < 0.05; ** = P-value < 0.01.

While the rest of our study focused on coding variants, *i.e*. variants that likely have a direct effect on protein function, we additionally aimed to get insight into the impact of non-coding common variants. Indeed, accumulating evidence highlights a role for common non-coding genetic variation in human health(46, 47), in inflammatory responses(52, 53), and even specifically in innate immune responses(54–56). Therefore, for the purpose of replicating our coding common variant signals, we repeated our significantly associated common variant sets in this study with previously published genotyping data from the same (FG500) cohort containing coding and non-coding common genome-wide genetic variation(31). All common variants (cohort AF>0.05) within a set were pruned for Linkage Disequilibrium (LD), using within cohort R^2^ metrics. **Figure 5B** shows that by subjecting these expanded sets to the same SKAT, we identified associations between *C. albicans-induced* IL-6 production and non-coding common variants in *IL36A* (P-value=0.049), *IL38* (P-value=0.007) and pro-inflammatory genes (P-value=0.019), and between *C. albicans-induced* IL-1ß production and non-coding common variants in *IL38* (P-value=0.046). We visualized these associations by calculating an allelic score, with single SNP directions and MAF-based variant weights incorporated, for all significant sets in **5B**. In **Figure 5C** the weighted directional allelic score is displayed in correlation with cytokine production, demonstrating that for each set there is an increasing cytokine production with increasing score, highest for *IL38* set with a correlation (R^2^) of 0.025. Finally, to evaluate the individual contribution of non-coding common variants in a set, we organized single SNP effect estimates based on their direction and significance (**Additional File 7**). A more in-depth follow up on *IL36A* set non-coding SNP rs1562305, revealed its non-coding activity by a combination of an interaction with *IL36G* and *IL36A* based on K562 Hi-C data and Pol II-associated chromosomal contacts are observed in the regions surrounding rs1562305 and *IL36A* (**Additional File 8A and B**). Moreover, rs1562305 falls within a region that is bound by several transcription factors, including *HNF4A*, that has been associated with ulcerative colitis and diabetes mellitus type II (**Additional File 8C**)(57).

## DISCUSSION

In this study we identify and characterize rare and common genetic variation in genes related to the IL-1 pathway, and determine their impact on the inter-individual variability of stimulus-specific *in vitro* cytokine responses, measured in whole blood from healthy individuals. In addition, we assess the relative contribution of rare as compared to common variants, as well as the joint effect of rare and common variants, by employing various grouping strategies.

Over the past decades, multiple studies have identified a role for common genetic variation on cytokine level and response, however a significant proportion of inter-individual variability remains to be determined(17–20). Common and rare variants have mostly been studied separately, and the differentiation between them is an arbitrary decision. Considering increasing evidence that specific combinations of variants with variable frequencies accounts for variability in phenotypic presentation(12–14), in particular for a combination of phenotypic characteristics that do not fit one specific clinical diagnosis(58), we hypothesized this concept might also apply to the inter-individual variability in cytokine responses. As such, this study aimed to provide a more holistic view, that considers the interplay of variants of different allele frequencies. For this purpose, we sequenced the coding regions of 48 genes related to the IL-1 pathway in almost 500 healthy individuals, and assessed *in vitro* IL-1ß and IL-6 production by whole blood in response to LPS, PHA, *C. albicans* and *S. aureus*. By means of the Sequence Kernel Association Test (SKAT), we tested for association between cytokine production and only common- (SKAToC), only rare- (SKAToR), and common and rare variants combined (SKATjoint), over various levels of grouping strategies; gene-, subpathway, and inflammatory-phenotype groups.

The strongest rare variant association detected in this study was between *CASP1* and LPS-induced IL-6 cytokine production. CASP-1 protein, encoded by *CASP1* gene, is responsible for cleavage of the inactive mediators IL-1ß, IL-18 and IL-33 into their active form. The association in this study was based on five rare variants: two private variants (one new to public databases); two variants identified in two individuals in heterozygous state, and one variant identified in five individuals in homozygous state (see **Additional File 5**). The fact that we observed a burden of rare variants not with IL-1ß but with IL-6 production, may suggest an unknown effect of CASP-1 on IL-6 production and/or release, although this is a speculation and remains to be demonstrated in future studies. Alternatively, immune responses are highly dynamic and as such can be influenced differentially by genetics over time(59), and since we used cytokine measurements after 48 hours of stimulation, it could also reflect an unnoticed effect of these variants on preceding IL-1ß production, that subsequently influence the induction of IL-6.

We detected an almost equally strong rare variant burden between *NCF4* and both IL-1ß and IL-6 production in response to PHA stimulation. This association was based on three private rare variants, one of which is new to public databases. Two variants are located in the canonical splice-acceptor sites, thereby possibly affecting the splicing of this gene, which most likely leads to loss-of-function variants with reduced RNA and possibly protein levels as a consequence. The *NCF4* gene encodes the NCF4 protein which is part of the cytoplasmic unit of the NADPH (nicotinamide adenine dinucleotide phosphate) oxidase enzyme system involved in phagocytosis(60). Variation in NADPH related genes are clinically relevant for human immunity as, rare mutations have been linked to chronic granulomatous disease (CGD, a primary immunodeficiency), and common variants have been associated with immune related diseases like Crohn’s disease and systemic lupus erythematodes (SLE)(61, 62). Knock-out of *Ncf4* in mice results in a defective NADPH-complex, with as a consequence reduced ROS-production and inefficient phagocytosis, ultimately leading to increased cytokine production(63, 64). The association detected in this study between *NCF4* rare variants and decreased cytokine production therefore suggests a gain-of-function effect on the NADPH-complex. However, considering the splice-affecting characteristics of two of these rare variants, the resulting effect on the NADPH oxidase system remains inconclusive and requires further investigation.

In addition, our study revealed an association between rare variants in anti-inflammatory genes and *S. aureus*-induced IL-6 production. The unidirectional characteristic of this was reflected by the observation that in whole blood from individuals carrying a rare variant more IL-6 cytokine was produced in response to *S. aureus*, as confirmed by Wilcoxon rank-sum P-value=0.0054 (**Figure 3C**). Remarkably, more than half of the anti-inflammatory genes are autophagy genes, which is in line with the notion that defective autophagy results in increased cytokine production, with increased inflammatory disease severity, like colitis and CGD, as a consequence(63, 65).

An intrinsic issue with rare variants is their low-frequency, resulting in limited power for association testing(66). Combining multiple rare variants followed by appropriate combination tests can improve power, but requires prior knowledge on the biological effects of the genes studied. Our IL-1 subpathway rare variant association with LPS-induced IL-1ß production confirms that appropriate grouping produces a significant association (SKAToR _adj_P-value=0.003), even though the individual genes did not (**Figure 2A**). The most likely reason for this is insufficient power, and indeed a thorough inspection of the raw data disclosed significant underlying gene-level association P-values that did not survive the Bonferroni-adjustment. The subpathway level association between common variants in genes of the IL-30s subpathway and IL-6 production in response to *C. albicans*, illustrates that even for common variants appropriate grouping can increase power. For instance, the IL-30s subpathway association with *C. albicans* stimulated IL-6 cytokine we detected, was more significant than could be explained by the gene-level associations with *IL36A* and *IL38*. Even though common variants in other members of the respective subpathway (*i.e. SIGIRR* and *IL37*) individually were not associated, most likely they do contribute to the subpathway signal. The *C. albicans-specific* common variant signature identified in this study by combining multiple common variants in a set over several levels of magnitude, additionally supports Smeekens et al.(67), where they identified the innate immune pathway to be induced by *C. albicans*. Next to highlighting the role for coding common variants, our validation with non-coding common variants in the same cohort supports the importance of considering interplay between multiple common variants, and additionally substantiates the urge for studying the impact of epigenetic regulation in immunity(68). Moreover, we identified a potential epigenetic mechanism that may allow insights into the gene regulation by non-coding variants, for rs1562305 on *IL36A* and *IL36G* gene expression levels (**Additional File 8**).

Another method to increase power in rare variant association studies, is to use extreme phenotypes for the purpose of enriching rare variants with strong(er) effects in outlier individuals(24, 44, 45). In this study, we used 1% extreme cytokine producers in a binary association and overlapped the results with our continuous tests, for the purpose of 1) characterization of stimulus-specific mechanisms, and 2) providing an indication of direction. Our strongest continuous rare variant associations (*CASP1* with LPS-induced IL-6 cytokine and *NCF4* with PHA-induced IL-1ß and IL-6 cytokine), reemerged in low-responders. The fact that these associations were identified both in our continuous and binary analyses, is in line with the finding that extreme phenotyping can enrich for the presence of rare causal variants(44, 45). Remarkably, our *NCF4* private variant carriers were restricted to the 1% lowest PHA-induced IL-6 cytokine response, substantiating that most extreme variants (in terms of lowest frequency) can have most extreme effects on the phenotype level(39, 69–71). In addition, we identified recurrent associations between LPS-induced IL-1ß cytokine production and five genes of the IL-1 subpathway, substantiating the subpathway level association and at the same time reflecting lack of power in the identification of individual gene associations. Interestingly, we identified an association between *S. aureus*-induced IL-1ß cytokine production and rare variants in *IL18BP*, recurrent in the top-responders. The encoded protein (IL-18BP) exerts major anti-inflammatory effects by inhibiting IL-18 signaling(72), and previous studies have shown that IL-18BP levels are negatively correlated with cytokine production by lymphocytes(17). The identified burden in top-responders here, suggests that rare variants in *IL18BP* negatively affect IL-18BP levels, allowing for higher *S. aureus*-induced cytokine production. The role of IL-18/IL-18BP pathway in cytokine responses to *Staphylococci* therefore needs further investigation, as it has the potential to provide important new insights in severe diseases caused by this pathogenic microorganism (*e.g*. sepsis).

Our study cohort is one of the largest to date in which extensive immunophenotyping experiments have been performed(17, 18, 20). The associations described here are based on cytokine production by whole blood, as it is most comparable to the *in vivo* situation and therefore most representative for physiologic immune responses. Furthermore, the investigated stimuli were chosen as representatives for an array of microbial infections, specifically: LPS is expressed on the bacterial cell wall of Gram-negative bacteria; PHA is synthesized by *Bacillus Rhodococcus* and *Pseudomonas species*; and *C. albicans* and *S. aureus* are major invading pathogens representative of fungi and Gram-positive bacteria, respectively. Future efforts investigating a broader array of pathogens, as well as specific contribution of immune cell subtypes, based on the additional publicly available data for this cohort(31) is therefore highly recommended. Nevertheless, potential limitations of this study cohort include the relatively small sample size and cohort characteristics (restricted age distribution and residency), and replication in a larger cohort for validation is favorable. Secondly, despite the cost-effectiveness of MIP-sequencing (*e.g*. ± €25,-per sample for the IL-1 panel), larger intronic or non-coding regions are not sequenced and as such escape analysis. The potential of using whole genome sequencing data to investigate the role of rare non-coding genetic variation, thereupon seems promising. Thirdly, the SKAT is powerful, but computes only set-wise association P-values and does not provide single variant effect estimates, neither does it provide direction in terms of positive/negative effects or increased/decreased risk. As we cannot exclude interaction effects between variants in a set, the contribution of single variants to a phenotype is difficult to estimate and as such the clinical applicability remains complex and requires more in-depth follow-up. Lastly, this study was designed for the purpose of discovering correlations between genetic variants and functional cytokine read-outs, but with the prospect of future, in-depth functional follow-up studies. In order to maximize the discovery layer, we Bonferroni-adjusted p-values based on the number of groups tested per stimulus-cytokine combination, frequency and grouping-level, and as such we did not apply post-hoc corrections.

## CONCLUSIONS

In conclusion, this study shows that common and rare genetic variation in genes of the IL-1 pathway, separately and combined, differentially influence *in vitro* IL-1ß and IL-6 cytokine responses induced by various stimuli. Not only do we add to the knowledge on the role of common variants in the IL-1 pathway, we additionally highlight the important role of rare variants (alone or in combination) on immune response variability. On a broader perspective, this study provides insight into potential mechanisms that are translatable to new hypothesis-driven identification of rare variant involvement in inflammatory and immunological mechanisms and diseases. Over and above, the framework used in this study is expandable to a wide variety of (non-immune) complex phenotypes, and as such can lead the way to new insights and theories for any phenotype of interest, and therefore has the potential to contribute to better understanding of unresolved, complex diseases.

## Supporting information

Additional File 1. IL-1 pathway grouping strategies.

Additional File 2. Baseline characteristics.

Additional File 3. MIPs covering 48 IL-1 pathway genes.

Additional File 4. Average MIP-coverage.

Additional File 5. Complete variant list.

Additional File 6. All SKAT output.

Additional File 7. Non-coding SNP set effect estimates.

Additional File 8. Epigenetic effects of rs1562305.

## ADDITIONAL FILES

*Additional File 1*. Variant grouping strategies of 48 Interleukin-1 pathway related genes.

*Additional File 2*. Baseline characteristics of healthy individuals (n=463).

*Additional File 3*. Molecular Inversion Probes (MIPs) covering all coding exons of 48 genes of the Interleukin-1 pathway.

*Additional File 4*. Average coverage depth per gene and overall for healthy individuals (n=463) included in analysis.

*Additional File 5*. Complete variant list.

*Additional File 6*. All SKAT output.

*Additional File 7*. Non-coding SNP set effect estimates distribution.

*Additional File 8*. Epigenetic effects of IL36A set non-coding SNP rs1562305.

## ABBREVIATIONS

_adj_P-value: Bonferroni-adjusted P-value
AF: Allele frequency
AoSD: Adult-onset Still’s Disease
BAQ: Base Alignment Quality
BT: Binary trait
BWA-MEM: Burrows-Wheeler Aligner
*C. albicans*: *Candida albicans*
CADD: Combined Annotation Dependent Depletion
CAPS: Cryopyrin associated periodic syndromes
CFU: Colony-forming unit
CGD: Chronic granulomatous disease
ChiA-PET: Chromatin interaction analysis with paired-end tag
ChiP-seq: Chromatin immunoprecipitation sequencing
CT: Continuous trait
dbSnp: Single Nucleotide Polymorphism Database
DIRA: DIRA: Deficiency of IL-1 receptor antagonist
DNA: Desoxyribonucleic acid
*e.g*.: *exempli gratia*
ELISA: Enzyme-linked immunosorbent assay
ExAc: Exome Aggregation Consortium
FG: Functional Genomics
GATK: Genome Analysis Toolkit
gnomAD: Genome Aggregation Database
GWAS: Genome Wide Association Studies
HFGP: Human Funtional Genomics Project
*i.e*.: *id est*
IL-1β: Interleukin-1β
IL-6: Interleukin-6
IL: Interleukin
kB: Kilobase
LD: Linkage disequilibrium
_LOW_BT: Binary trait association analysis using 1% lowest cytokine producers
LPS: Lipopolysaccharide
MAF: Minor allele frequency
MAS: Macrophage activation syndrome
MIP: Molecular Inversion Probe
μg: Microgram
μL: Microliter
mL: Milliliter
NADPH: Nicotinamide adenine dinucleotide phosphate
NCBI RefSeq: National Center for Biotechnology Information Reference Sequence
ng: Nanogram
PHA: Phytohaemagglutinin
PMNs: Polymorphonuclear cells
QUAL: Quality parameter in vcf
R^2^: Correlation metric
RA: Rheumatoid arthritis
Radboudumc: Radboud university medical center
RNA: Ribonucleic acid
ROS: Reactive Oxygen Species
RVBA: Rare variant burden analysis
*S. aureus:*: *Staphylococcus aureus*
SKAT: Sequence Kernel Association Test
SKATjoint: SKAT common and rare variants
SKATO/SKAT-O: linear combination of the Burden Test and SKAT with optimal weights
SKAToC: SKAT only common variants
SKAToR: SKAT only rare variants
SLE: Systemic lupus erythematodes
SNP: Single Nucleotide Polymorphism
tagSnps: Set of LD-pruned SNPs
_TOP_BT: Binary trait association analysis using 1% highest cytokine producers
vcf: Variant Call Format
VEP: Variant Effect Predictor

## ACKNOWLEDGEMENTS

The authors thank the Radboud Genomics Technology Center, Radboud University Medical Center (Radboudumc), Nijmegen for support in sequencing. LABJ was supported by a Competitiveness Operational Programme grant of the Romanian Ministry of European Funds (HINT, P_37_762, MySMIS 103587). CAD was supported by NIH AI-15614. MMM was supported by the South African Medical Research Council, Department of Science & Technology of South Africa, Chan Zuckerberg Initiative and Bill & Melinda Gates Foundation. VK was supported by Hypatia tenure track fellowship. MGN was supported by the Nederlandse Organisatie voor Wetenschappelijk onderzoek (Spinoza grant), and the European Research Council (grant agreement No 310372). FLV and AH were supported by the Interleukin Foundation. AH was supported by the SOLVE-RD project, funded by the European Union’s Horizon 2020 research and innovation programme under grant agreement No 779257.

## AUTHOR CONTRIBUTIONS

MGN, FLvdV and AH designed the project. RCvD, MGN, AH conceptualized the experiments and analysis. PA, MJ, MS performed the experiments. RCvD performed data analysis and data visualization. MvdV and CG supported the data analysis. MMM performed additional data analysis. GC, VK, CAD, LABJ, FLvdV and MGN helped with data interpretation. RCvD, MGN and AH wrote the manuscript. All authors contributed to and approved the final manuscript.

## DATA AVAILABILITY

All demographic-, immunophenotyping- and genotyping-data from the FG500-cohort used in this study is publicly available on the BBMRI-NL archive(31). All variants called in MIP-sequencing data based on the IL-1 panel from the same cohort are published here, as well as all association results.

## CODE AVAILABILITY

Code for processing and filtering MIP-based sequencing data are extensively explained in the methods section of this manuscript and will be made available upon reasonable request. The source code from the R packages used in this study are freely available online. Code for processing the output and generating the figures will likewise be made available upon reasonable request.

## References

1. Parham P, Janeway CI. The immune system. 3rd ed. ed. London: Garland Science; 2009.

2. Dinarello CA. Overview of the IL-1 family in innate inflammation and acquired immunity. Immunol Rev. 2018;281(1):8–27.

3. Garlanda C, Dinarello CA, Mantovani A. The interleukin-1 family: back to the future. Immunity. 2013;39(6):1003–18.

4. Sun Q, Fan J, Billiar TR, Scott MJ. Inflammasome and autophagy regulation - a two-way street. Mol Med. 2017;23:188–95.

5. Dinarello CA. The IL-1 family of cytokines and receptors in rheumatic diseases. Nat Rev Rheumatol. 2019;15(10):612–32.

6. Palomo J, Dietrich D, Martin P, Palmer G, Gabay C. The interleukin (IL)-1 cytokine family--Balance between agonists and antagonists in inflammatory diseases. Cytokine. 2015;76(1):25–37.

7. Jesus AA, Goldbach-Mansky R. IL-1 blockade in autoinflammatory syndromes. Annu Rev Med. 2014;65:223–44.

8. Wjst M, Sargurupremraj M, Arnold M. Genome-wide association studies in asthma: what they really told us about pathogenesis. Curr Opin Allergy Clin Immunol. 2013;13(1):112–8.

9. Korver SK, Gibson RJ, Bowen JM, Coller JK. Toll-like receptor/interleukin-1 domain innate immune signalling pathway genetic variants are candidate predictors for severe gastrointestinal toxicity risk following 5-fluorouracil-based chemotherapy. Cancer Chemother Pharmacol. 2019;83(2):217–36.

10. Tekola Ayele F, Doumatey A, Huang H, Zhou J, Charles B, Erdos M, et al. Genome-wide associated loci influencing interleukin (IL)-10, IL-1Ra, and IL-6 levels in African Americans. Immunogenetics. 2012;64(5):351–9.

11. He M, Cornelis MC, Kraft P, van Dam RM, Sun Q, Laurie CC, et al. Genome-wide association study identifies variants at the IL18-BCO2 locus associated with interleukin-18 levels. Arterioscler Thromb Vasc Biol. 2010;30(4):885–90.

12. Chung RH, Kang CY. A Powerful Gene-Based Test Accommodating Common and Low-Frequency Variants to Detect Both Main Effects and Gene-Gene Interaction Effects in Case-Control Studies. Front Genet. 2017;8:228.

13. Hsieh AR, Chen DP, Chattopadhyay AS, Li YJ, Chang CC, Fann CSJ. A non-threshold region-specific method for detecting rare variants in complex diseases. PLoS One. 2017;12(11):e0188566.

14. Ionita-Laza I, Lee S, Makarov V, Buxbaum JD, Lin X. Sequence kernel association tests for the combined effect of rare and common variants. Am J Hum Genet. 2013;92(6):841–53.

15. Wagih O, Galardini M, Busby BP, Memon D, Typas A, Beltrao P. A resource of variant effect predictions of single nucleotide variants in model organisms. Mol Syst Biol. 2018;14(12):e8430.

16. Boyle EA, Li YI, Pritchard JK. An Expanded View of Complex Traits: From Polygenic to Omnigenic. Cell. 2017;169(7):1177–86.

17. Bakker OB, Aguirre-Gamboa R, Sanna S, Oosting M, Smeekens SP, Jaeger M, et al. Integration of multi-omics data and deep phenotyping enables prediction of cytokine responses. Nat Immunol. 2018;19(7):776–86.

18. Li Y, Oosting M, Smeekens SP, Jaeger M, Aguirre-Gamboa R, Le KTT, et al. A Functional Genomics Approach to Understand Variation in Cytokine Production in Humans. Cell. 2016;167(4):1099–110 e14.

19. Piasecka B, Duffy D, Urrutia A, Quach H, Patin E, Posseme C, et al. Distinctive roles of age, sex, and genetics in shaping transcriptional variation of human immune responses to microbial challenges. Proc Natl Acad Sci U S A. 2018;115(3):E488–E97.

20. Ter Horst R, Jaeger M, Smeekens SP, Oosting M, Swertz MA, Li Y, et al. Host and Environmental Factors Influencing Individual Human Cytokine Responses. Cell. 2016;167(4):1111–24 e13.

21. Kim S, Becker J, Bechheim M, Kaiser V, Noursadeghi M, Fricker N, et al. Characterizing the genetic basis of innate immune response in TLR4-activated human monocytes. Nat Commun. 2014;5:5236.

22. Scepanovic P, Alanio C, Hammer C, Hodel F, Bergstedt J, Patin E, et al. Human genetic variants and age are the strongest predictors of humoral immune responses to common pathogens and vaccines. Genome Med. 2018;10(1):59.

23. Asimit J, Zeggini E. Testing for rare variant associations in complex diseases. Genome Med. 2009;1(11):24.

24. Auer PL, Lettre G. Rare variant association studies: considerations, challenges and opportunities. Genome Med. 2015;7(1):16.

25. Bodmer W, Bonilla C. Common and rare variants in multifactorial susceptibility to common diseases. Nat Genet. 2008;40(6):695–701.

26. Hiatt JB, Pritchard CC, Salipante SJ, O’Roak BJ, Shendure J. Single molecule molecular inversion probes for targeted, high-accuracy detection of low-frequency variation. Genome Res. 2013;23(5):843–54.

27. Jansen S, Hoischen A, Coe BP, Carvill GL, Van Esch H, Bosch DGM, et al. A genotype-first approach identifies an intellectual disability-overweight syndrome caused by PHIP haploinsufficiency. Eur J Hum Genet. 2018;26(1):54–63.

28. Neveling K, Mensenkamp AR, Derks R, Kwint M, Ouchene H, Steehouwer M, et al. BRCA Testing by Single-Molecule Molecular Inversion Probes. Clin Chem. 2017;63(2):503–12.

29. O’Roak BJ, Vives L, Fu W, Egertson JD, Stanaway IB, Phelps IG, et al. Multiplex targeted sequencing identifies recurrently mutated genes in autism spectrum disorders. Science. 2012;338(6114):1619–22.

30. Human Functional Genomics Project Home Site. http://www.humanfunctionalgenomics.org/site/. Accessed 8 April 2019.

31. Human Functional Genomics Project BBMRI-NL archive. https://hfgp.bbmri.nl/. Accessed 8 April 2019.

32. Boyle EA, O’Roak BJ, Martin BK, Kumar A, Shendure J. MIPgen: optimized modeling and design of molecular inversion probes for targeted resequencing. Bioinformatics. 2014;30(18):2670–2.

33. Li H, Durbin R. Fast and accurate short read alignment with Burrows-Wheeler transform. Bioinformatics. 2009;25(14):1754–60.

34. Van der Auwera GA, Carneiro MO, Hartl C, Poplin R, Del Angel G, Levy-Moonshine A, et al. From FastQ data to high confidence variant calls: the Genome Analysis Toolkit best practices pipeline. Curr Protoc Bioinformatics. 2013;43:11 0 1- 0 33.

35. Smigielski EM, Sirotkin K, Ward M, Sherry ST. dbSNP: a database of single nucleotide polymorphisms. Nucleic Acids Res. 2000;28(1):352–5.

36. Li H, Handsaker B, Wysoker A, Fennell T, Ruan J, Homer N, et al. The Sequence Alignment/Map format and SAMtools. Bioinformatics. 2009;25(16):2078–9.

37. McLaren W, Gil L, Hunt SE, Riat HS, Ritchie GR, Thormann A, et al. The Ensembl Variant Effect Predictor. Genome Biol. 2016;17(1):122.

38. Kircher M, Witten DM, Jain P, O’Roak BJ, Cooper GM, Shendure J. A general framework for estimating the relative pathogenicity of human genetic variants. Nat Genet. 2014;46(3):310–5.

39. Lek M, Karczewski KJ, Minikel EV, Samocha KE, Banks E, Fennell T, et al. Analysis of protein-coding genetic variation in 60,706 humans. Nature. 2016;536(7616):285–91.

40. Wu MC, Lee S, Cai T, Li Y, Boehnke M, Lin X. Rare-variant association testing for sequencing data with the sequence kernel association test. Am J Hum Genet. 2011;89(1):82–93.

41. Lee S, Emond MJ, Bamshad MJ, Barnes KC, Rieder MJ, Nickerson DA, et al. Optimal unified approach for rare-variant association testing with application to small-sample case-control whole-exome sequencing studies. Am J Hum Genet. 2012;91(2):224–37.

42. Hernandez RD, Uricchio LH, Hartman K, Ye C, Dahl A, Zaitlen N. Ultrarare variants drive substantial cis heritability of human gene expression. Nat Genet. 2019;51(9): 1349–55.

43. Li X, Kim Y, Tsang EK, Davis JR, Damani FN, Chiang C, et al. The impact of rare variation on gene expression across tissues. Nature. 2017;550(7675):239–43.

44. Barnett IJ, Lee S, Lin X. Detecting rare variant effects using extreme phenotype sampling in sequencing association studies. Genet Epidemiol. 2013;37(2):142–51.

45. Bjornland T, Bye A, Ryeng E, Wisloff U, Langaas M. Powerful extreme phenotype sampling designs and score tests for genetic association studies. Stat Med. 2018;37(28):4234–51.

46. Gorkin DU, Qiu Y, Hu M, Fletez-Brant K, Liu T, Schmitt AD, et al. Common DNA sequence variation influences 3-dimensional conformation of the human genome. Genome Biol. 2019;20(1):255.

47. Quinn JP, Savage AL, Bubb VJ. Non-coding genetic variation shaping mental health. Curr Opin Psychol. 2019;27:18–24.

48. Li Y, Oosting M, Deelen P, Ricano-Ponce I, Smeekens S, Jaeger M, et al. Inter-individual variability and genetic influences on cytokine responses to bacteria and fungi. Nat Med. 2016;22(8):952–60.

49. Võsa U, Claringbould A, Westra H-J, Bonder MJ, Deelen P, Zeng B, et al. Unraveling the polygenic architecture of complex traits using blood eQTL metaanalysis. bioRxiv. 2018:447367.

50. Burgess S, Thompson SG. Use of allele scores as instrumental variables for Mendelian randomization. Int J Epidemiol. 2013;42(4):1134–44.

51. Rao SS, Huntley MH, Durand NC, Stamenova EK, Bochkov ID, Robinson JT, et al. A 3D map of the human genome at kilobase resolution reveals principles of chromatin looping. Cell. 2014;159(7):1665–80.

52. Meddens CA, van der List ACJ, Nieuwenhuis EES, Mokry M. Non-coding DNA in IBD: from sequence variation in DNA regulatory elements to novel therapeutic potential. Gut. 2019;68(5):928–41.

53. Ramsuran V, Ewy R, Nguyen H, Kulkarni S. Variation in the Untranslated Genome and Susceptibility to Infections. Front Immunol. 2018;9:2046.

54. Zhang Q, Chao TC, Patil VS, Qin Y, Tiwari SK, Chiou J, et al. The long noncoding RNA ROCKI regulates inflammatory gene expression. EMBO J. 2019;38(8).

55. Blecher-Gonen R, Amit I. M(odu)LLating the innate response. Immunity. 2012;36(4):551–2.

56. Austenaa L, Barozzi I, Chronowska A, Termanini A, Ostuni R, Prosperini E, et al. The histone methyltransferase Wbp7 controls macrophage function through GPI glycolipid anchor synthesis. Immunity. 2012;36(4):572–85.

57. Lees CW, Barrett JC, Parkes M, Satsangi J. New IBD genetics: common pathways with other diseases. Gut. 2011;60(12):1739–53.

58. Tcheandjieu C, Aguirre M, Saha P, Potiny P, Haendel M, Ingelsson E, et al. A phenome-wide association study of four syndromic genes reveals pleiotropic effects of common and rare variants in the general population. medRxiv. 2019:19010736.

59. Kim-Hellmuth S, Bechheim M, Putz B, Mohammadi P, Nedelec Y, Giangreco N, et al. Genetic regulatory effects modified by immune activation contribute to autoimmune disease associations. Nat Commun. 2017;8(1):266.

60. Tarazona-Santos E, Machado M, Magalhaes WC, Chen R, Lyon F, Burdett L, et al. Evolutionary dynamics of the human NADPH oxidase genes CYBB, CYBA, NCF2, and NCF4: functional implications. Mol Biol Evol. 2013;30(9):2157–67.

61. Jacob CO, Eisenstein M, Dinauer MC, Ming W, Liu Q, John S, et al. Lupus-associated causal mutation in neutrophil cytosolic factor 2 (NCF2) brings unique insights to the structure and function of NADPH oxidase. Proc Natl Acad Sci U S A. 2012;109(2):E59–67.

62. Rioux JD, Xavier RJ, Taylor KD, Silverberg MS, Goyette P, Huett A, et al. Genome-wide association study identifies new susceptibility loci for Crohn disease and implicates autophagy in disease pathogenesis. Nat Genet. 2007;39(5):596–604.

63. de Luca A, Smeekens SP, Casagrande A, Iannitti R, Conway KL, Gresnigt MS, et al. IL-1 receptor blockade restores autophagy and reduces inflammation in chronic granulomatous disease in mice and in humans. Proc Natl Acad Sci U S A. 2014;111(9):3526–31.

64. Winter S, Hultqvist Hopkins M, Laulund F, Holmdahl R. A Reduction in Intracellular Reactive Oxygen Species Due to a Mutation in NCF4 Promotes Autoimmune Arthritis in Mice. Antioxid Redox Signal. 2016;25(18):983–96.

65. Cavalli G, Cenci S. Autophagy and Protein Secretion. J Mol Biol. 2020.

66. Sun J, Zheng Y, Hsu L. A unified mixed-effects model for rare-variant association in sequencing studies. Genet Epidemiol. 2013;37(4):334–44.

67. Smeekens SP, Ng A, Kumar V, Johnson MD, Plantinga TS, van Diemen C, et al. Functional genomics identifies type I interferon pathway as central for host defense against Candida albicans. Nat Commun. 2013;4:1342.

68. Winter DR, Jung S, Amit I. Making the case for chromatin profiling: a new tool to investigate the immune-regulatory landscape. Nat Rev Immunol. 2015;15(9):585–94.

69. Wainschtein P, Jain DP, Yengo L, Zheng Z, Cupples LA, Shadyab AH, et al. Recovery of trait heritability from whole genome sequence data. bioRxiv. 2019:588020.

70. de Magalhaes JP, Wang J. The fog of genetics: what is known, unknown and unknowable in the genetics of complex traits and diseases. EMBO Rep. 2019;20(11):e48054.

71. Kluck V, van Deuren RC, Cavalli G, Shaukat A, Arts P, Cleophas MC, et al. Rare genetic variants in interleukin-37 link this anti-inflammatory cytokine to the pathogenesis and treatment of gout. Ann Rheum Dis. 2020;79(4):536–44.

72. Novick D, Kim SH, Fantuzzi G, Reznikov LL, Dinarello CA, Rubinstein M. Interleukin-18 binding protein: a novel modulator of the Th1 cytokine response. Immunity. 1999;10(1):127–36.

