## Additional File 2. Baseline characteristics. for "Impact of rare and common genetic variation in the Interleukin-1 pathway on human cytokine responses"

**A.** Age distribution classified by gender

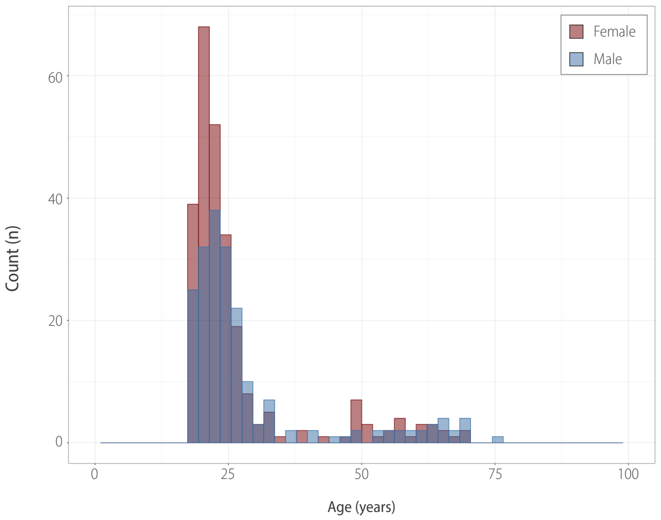


**B.** Cytokine distribution per stimulus classified by gender

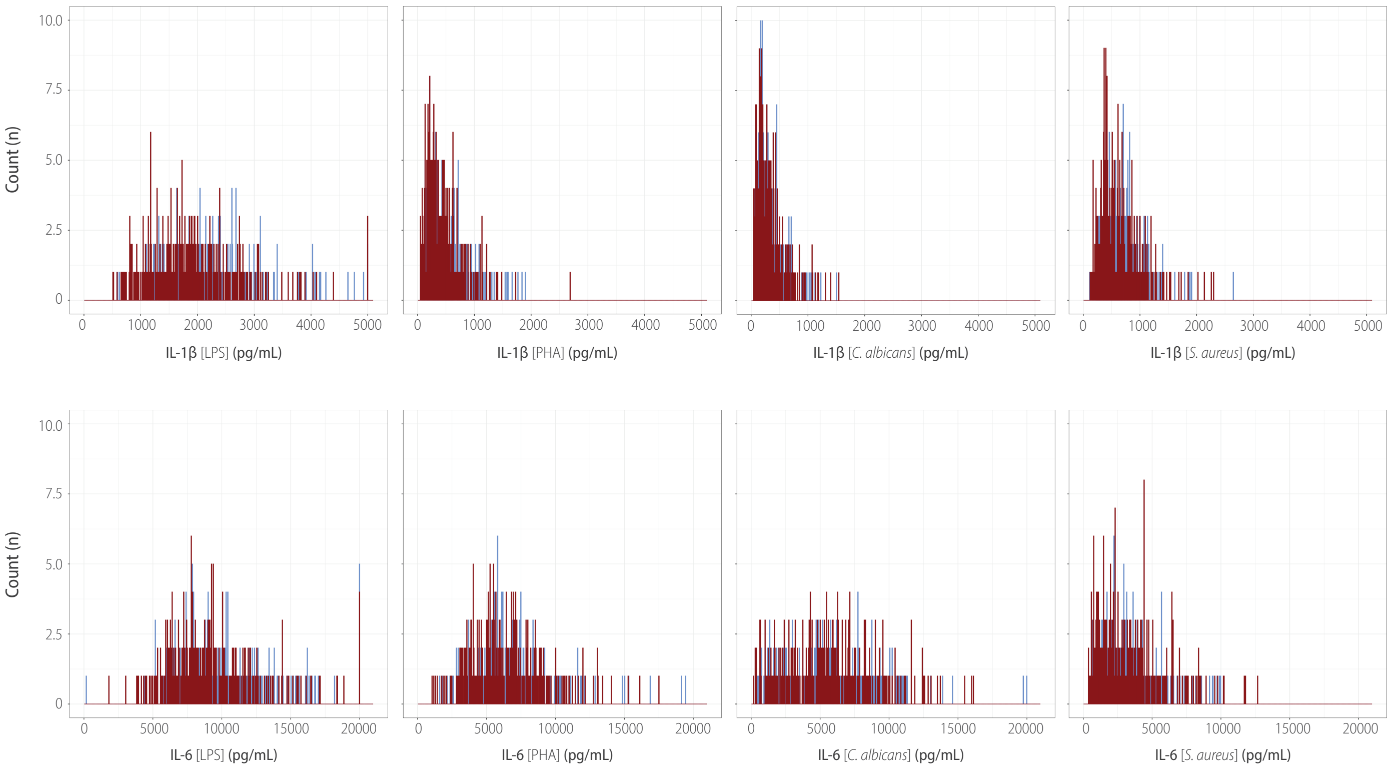


**Additional File 2. Baseline characteristics of healthy individuals (n=463). (A)** shows the age distribution of all individuals included in analyses classified by gender (mean age - Female=26.66, Male=28.84; median age - Female=22, Male=24). **(B)** shows the IL-1β and IL-6 *in vitro* cytokine production in response to LPS (100ng/mL), PHA (10µg/mL), *C. albicans* ($10^6$ CFU/mL), *S. aureus* ($1x10^6$/mL), classified by gender.
