## Additional File 4. Average MIP-coverage. for "Impact of rare and common genetic variation in the Interleukin-1 pathway on human cytokine responses"

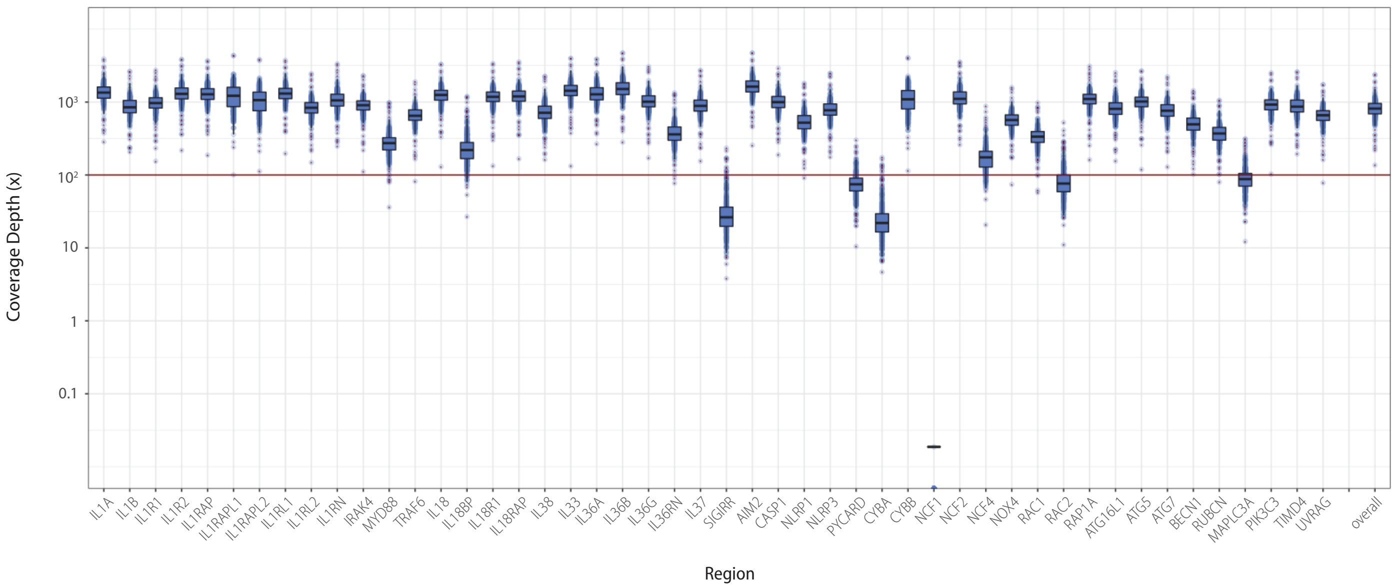


**Additional File 3. Average coverage depth per gene and overall for healthy individuals (n=463) included in analysis.** The average coverage for each individual per gene (and overall on the most right) is visualized in separate jitter plots, overlayed by boxplots, where boxes contain a median line, edges at 25$^{th}$ and 75$^{th}$ percentile, and default whiskers (1.5 * InterQuartileRange). Logarithmic y-axis, with a horizontal red line representing the analysis inclusion coverage cut-off of 100x.
