## Additional File 7. Non-coding SNP set effect estimates. for "Impact of rare and common genetic variation in the Interleukin-1 pathway on human cytokine responses"

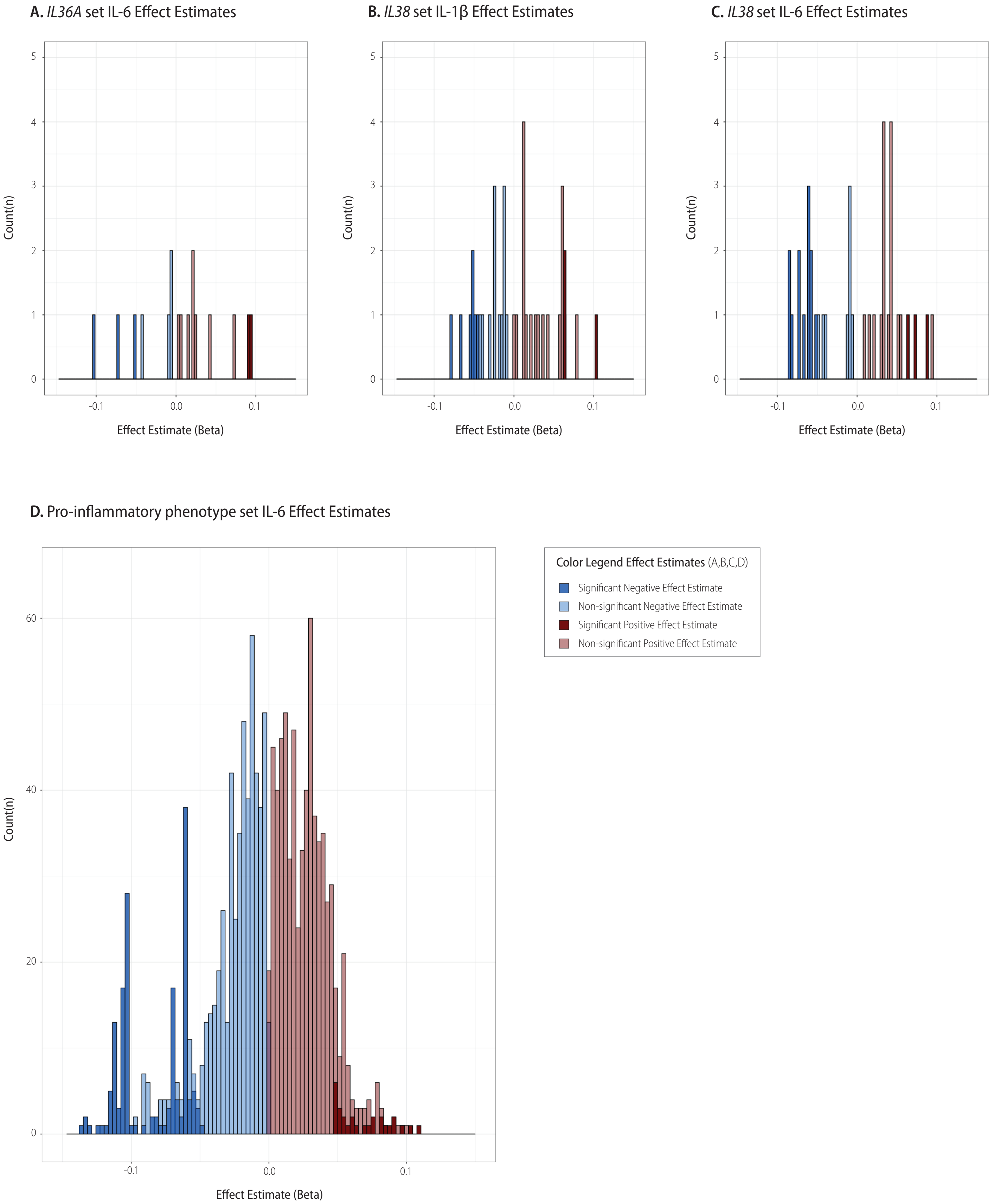


**Additional File 7. Non-coding SNP set Effect Estimate distribution.** The single SNP effect estimate distribution is visualized in a histogram for all non-coding SNPs in a set. Effect estimates were extracted from a linear model that was created for each SNP in a set using the residual *C. albicans* induced cytokine production as criterion variable and SNP as predictor variable.
