## Additional File 8. Epigenetic effects of rs1562305. for "Impact of rare and common genetic variation in the Interleukin-1 pathway on human cytokine responses"

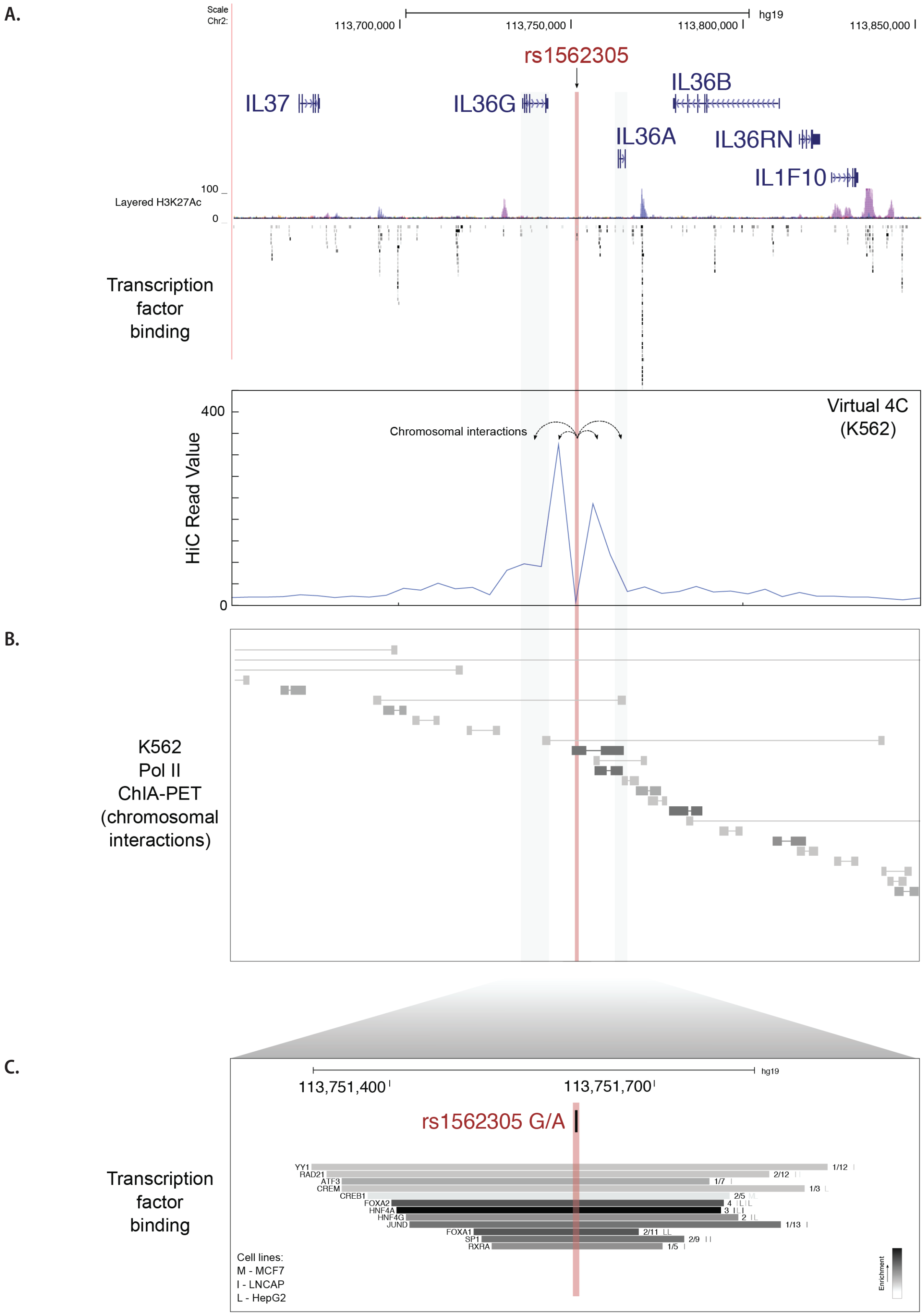


**Additional File 8. Epigenetic Effects of IL36A set non-coding SNP rs1562305. (A)** Visualization of K562 Hi-C data in virtual 4C format (in which rs1562305 is the viewpoint), predicts that rs1562305 interacts with *IL36G* and *IL36A*. **(B)** ChiA-PET data from a library constructed in K562s with an antibody that enriches for Pol II. Pol II-associated chromosomal contacts between the region surrounding rs1562305 and *IL36A* were observed. **(C)** Transcription factor ChIPseq analysis across 130 cell types revealed that rs1562305 falls within a region of DNA that is bound by several transcription factors, including *HNF4A*.
